# Temporal Requirements of SKN-1/NRF as a Regulator of Lifespan and Proteostasis in *Caenorhabditis elegans*

**DOI:** 10.1101/2020.11.24.395707

**Authors:** Danielle Grushko, Amir Levine, Hana Boocholez, Ehud Cohen

**Affiliations:** Department of Biochemistry and Molecular Biology, the Institute for Medical Research Israel – Canada (IMRIC), the Hebrew University School of Medicine, Jerusalem 91120, Israel

**Keywords:** C. elegans, Aging, Proteostasis, Insulin/IGF signaling, SKN-1/NRF

## Abstract

Lowering the activity of the Insulin/IGF-1 Signaling (IIS) cascade results in elevated stress resistance, enhanced protein homeostasis (proteostasis) and extended lifespan of worms, flies and mice. In the nematode *Caenorhabditis elegans* (*C. elegans*), the longevity phenotype that stems from IIS reduction is entirely dependent upon the activities of a subset of transcription factors including the Forkhead factor DAF-16/FOXO (DAF-16), Heat Shock Factor-1 (HSF-1), SKiNhead/Nrf (SKN-1) and ParaQuat Methylviologen responsive (PQM-1). While DAF-16 determines lifespan exclusively during early adulthood and governs proteostasis in early adulthood and midlife, HSF-1 executes these functions foremost during development. Despite the central roles of SKN-1 as a regulator of lifespan and proteostasis, the temporal requirements of this transcription factor were unknown. Here we employed conditional knockdown techniques and discovered that in *C. elegans*, SKN-1 is primarily important for longevity and proteostasis during late larval development through early adulthood. Our findings indicate that events that occur during late larval developmental through early adulthood affect lifespan and proteostasis and suggest that subsequent to HSF-1, SKN-1 sets the conditions, partially overlapping temporally with DAF-16, that enable IIS reduction to promote longevity and proteostasis. Our findings raise the intriguing possibility that HSF-1, SKN-1 and DAF-16 function in a coordinated and sequential manner to promote healthy aging.

## Introduction

For decades, aging was thought to be an entirely stochastic, uncontrolled process driven by the accumulation of cellular damage [1, 2]. This view has changed as it became evident that manipulating the activities of several genetic and metabolic pathways elevates stress resistance, enhances protein homeostasis (proteostasis) and extends lifespans of various organisms. Dietary restriction (DR) [3], reducing Insulin/IGF-1 signaling (IIS) [4], lowering the activity of the mitochondrial electron transport chain (ETC) [5] and of signaling that emanates from the reproductive system [6], were all found to slow the aging process. The IIS, probably the most prominent aging-regulating pathway, is a key regulator of development, stress resistance, metabolism and longevity of various organisms [4, 7–9].

In the nematode *Caenorhabditis elegans* (*C. elegans*), upon binding of one of its ligands, the lone insulin/IGF-1 receptor DAF-2 activates a signaling cascade which regulates the activity of a nexus of transcription factors through a highly conserved set of molecular components. DAF-2’s downstream kinases mediate the phosphorylation of the transcription factors DAF-16 [10, 11] and SKN-1 [12]. These phosphorylation events retain DAF-16 and SKN-1 in the cytosol, preventing them from regulating their target gene networks. Analogously, the IIS negatively regulates the activity of HSF-1 by preventing the phosphorylation of DDL-1, a protein that interacts with this transcription factor. Non-phosphorylated DDL-1 along with DDL-2 and HSB-1 form a complex of proteins that binds HSF-1 and retains it in the cytosol [13]. The IIS also governs the cellular localization of PQM-1, a transcription factor which responds to IIS reduction in opposition to DAF-16, and plays key roles in the IIS-controlled lifespan determining mechanism [14]. Thus, *daf-2* knockdown by either mutation or RNA interference (RNAi) hyper-activates HSF-1, DAF-16 and SKN-1, creating long-lived, youthful and stress-resistant worms. These longevity and stress resistance effects of *daf-2* knockdown are dependent upon each of the aforementioned transcription factors [12, 15, 16]. Similarly to worms, reduced IGF-1 signaling was shown to extend the lifespan of mice [9], and mutations in components of the same pathway correlate with extreme longevity of humans [7, 17], indicating that the aging-regulating roles of the IIS are conserved from worms to mammals.

### The alteration of aging protects worms and mammals from toxic protein aggregation

Maintaining the integrity of the proteome is vital for organismal functionality and viability. However, as an organism ages, its ability to maintain proteostasis declines [18, 19], enabling subsets of proteins to form potentially toxic aggregates that accrue within the cell [20]. In some cases, the accumulation of aggregated proteins underlies the development of a myriad of late-onset maladies including neurodegenerative disorders such as Alzheimer’s disease (AD) [21] and Huntington’s disease (HD) [22]. Aging is the major risk factor for the manifestation of neurodegeneration, a common feature in these late-onset diseases [23]. This raises the prospect that the alteration of aging could maintain proteostasis in the late stages of life thereby preventing, or at least delaying, the emergence of neurodegeneration. Indeed, IIS reduction [24–26], DR [27], ETC impairments [28] and germ cell ablation [29], were all found to promote proteostasis and protect model nematodes from toxic protein aggregation (proteotoxicity). These mechanistic links define proteostasis collapse as an inherent aspect of aging [30]. Importantly, all the aforementioned IIS regulated transcription factors; DAF-16, HSF-1 [24, 25], SKN-1[31] and PQM-1 [32] are involved in the regulation of proteostasis, raising the prospect that modulating the activities of these factors could extend healthspan through late stages of life. However, to maintain proteostasis and extend healthspan without affecting lifespan, it is critical to ascertain the temporal requirements of these factors as lifespan and proteostasis regulators.

### The temporal requirements of DAF-16 and HSF-1

Reducing the IIS at different stages of life via *daf-2* RNAi, identified that IIS reduction during reproductive adulthood (days 1-6) and no other stage of life, extends the lifespan of *C. elegans* [33]. Consistently, an increase in lifespan was observed in the fruit fly *Drosophila melanogaster* when dFOXO (the ortholog of DAF-16) was over-expressed during reproductive adulthood but not during any other stages of life.[8]. Surprisingly, we discovered that HSF-1 is of foremost importance for the determination of lifespan during the L2 larval stage, but also has a marginal effect on lifespan during reproductive adulthood [34]. DAF-16 and HSF-1 also exhibit distinct temporal requirements for proteostasis maintenance. While DAF-16 is dispensable for proteotoxicity protection during development and plays its counter-proteotoxic protective roles exclusively during adulthood, HSF-1 executes these functions mainly during development [35]. These distinct temporal patterns raise questions about the functional relationship between these two transcription factors, SKN-1 and the IIS.

Despite the central roles of SKN-1 as a regulator of lifespan downstream of the IIS [12] and via the DR pathway [36], as well as its influence on proteostasis [31], the temporal requirements of SKN-1 for these functions were unknown. To address this, we used the nematode *C. elegans* and a conditional RNAi knockdown technique and found that SKN-1 governs lifespan and proteostasis primarily during development and during early adulthood. These observations raise important questions regarding the functional relationship between DAF-16 and SKN-1 and raise the prospect that during development SKN-1 regulates the expression of genes that encode for DAF-16 co-factors that are needed to promote longevity and proteostasis in late stages of life.

## Materials and Methods

### Worm and RNAi strains

N_2_ (wild type, Bristol), *daf-2* (e1370), CL2006 (*unc-54*p::human Aβ_3-42_), CF512 (fer-15(b26)II; fem-1(hc17)IV) and ad1116 (*eat-2* mutant) worms were obtained from the Caenorhabditis Genetics Center (CGC, Minneapolis, MN), which is funded by the National Institutes of Health Office of Research Infrastructure Programs (P40 OD010440). AGD1246 worms were a generous gift of Dr. Andrew Dillin (University of California at Berkeley). AM140 and AM1126 worms were a generous gift of Dr. Richard Morimoto (Northwestern University). All worm strains were routinely grown at 15°C for maintenance. For experimentation all worms were kept at 20°C throughout life except for CF512 animals which are heat-sensitive feminized and were therefore grown at 25°C throughout development to prevent progeny, and then maintained at 20°C throughout adulthood. To reduce gene expression, we used bacterial strains expressing dsRNA: empty vector (pAD12), *skn-1* and *dcr-1* dsRNA expressing bacteria from the M. Vidal RNAi library. Each RNAi bacteria colony was grown at 37°C in LB with 100μg/ml ampicillin and then seeded on NG-ampicillin plates and supplemented with 100mM Isopropyl β-D-1-thiogalactopyranoside (IPTG 1mM final concentration).

### Expression analysis by quantitative real-time PCR (qRT-PCR)

Synchronized eggs were placed on NGM plates seeded with the indicated bacteria. The worms were grown from hatching until day one of adulthood unless otherwise indicated. The worm samples were then harvested and washed with M9 buffer to remove bacteria from the samples. Each worm pellet was re-suspended in 1M DTT and RA1 (solution from the NucleoSpin® RNA kit (Macherey-Nagel, Duren Germany #740955.50)) and frozen at −80°C overnight. After thawing the samples on ice, zirconium oxide beads (Next Advance, ZrB05) were added to the samples and the samples were homogenized at 4°C using a Bullet Blender® (Next Advance). To separate RNA from protein and other materials, samples underwent centrifugation at room temperature in a tabletop centrifuge. The NucleoSpin RNA isolation Kit (Macherey Nagel, Duren Germany #740955.50) was used according to the manufacturer instructions to extract RNA. cDNA was generated by reverse transcription of the total RNA samples with iScriptRT Advanced cDNA Synthesis Kit for RT-PCR (Bio-Rad, Hercules, CA; #170–8891;). qRT-PCR was performed in triplicates using the iTaqTM Universal SYBR® Supermix (Bio-Rad; #172–5124) and quantified in a CFX96TM Real-Time PCR Detection System (Bio-Rad). The levels were normalized to the levels of *act-1* and/or *pmp-3* cDNA.

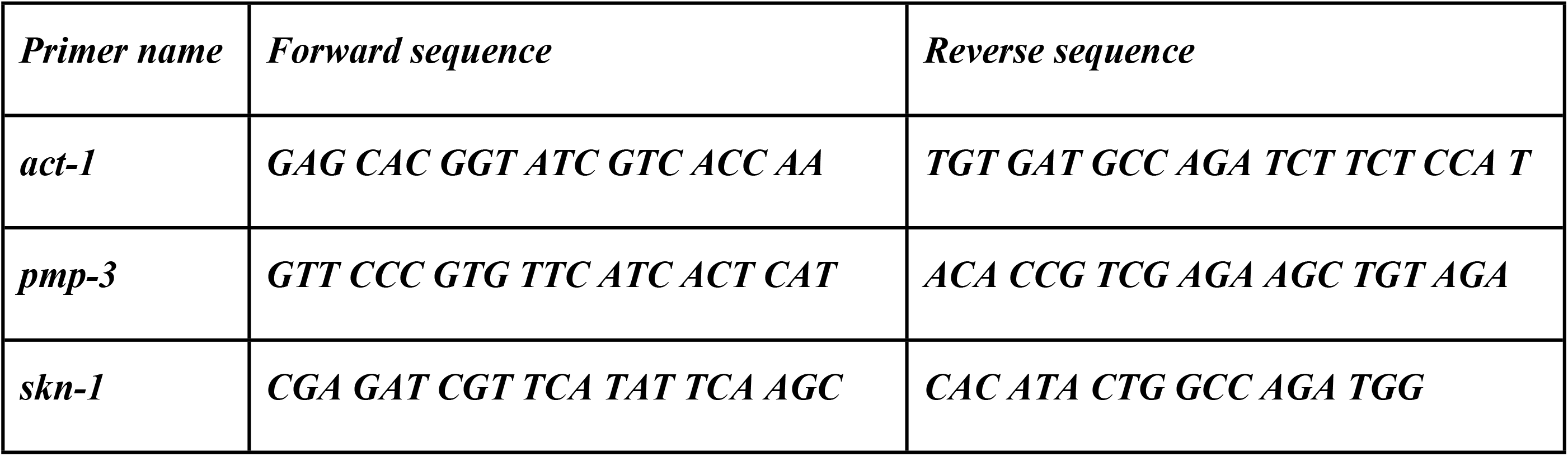

### Lifespan assay

Lifespan assays were conducted as previously described [34]. Briefly, synchronized eggs were placed on master NG-ampicillin 9cm plates seeded with the indicated RNAi bacterial strain and supplemented with 100mM IPTG (~1mM final concentration). CF512 worms were grown at 25°C throughout development to avoid progeny, then transferred to 20°C for the duration of their life. *daf-2* (e1370) mutant animals as well as N_2_ and ad1116 worms were developed and maintained at 20°C. At day 1 of adulthood, 120 animals per treatment were transferred onto 5cm NG-ampicillin plates (12 animals per plate). Worms that failed to move their heads when tapped twice with a platinum wire or when a hot pick was placed proximally to their body were scored as dead. Survival rates were recorded daily.

### Proteotoxicity assays

To follow Aβ-mediated toxicity by the “paralysis assay” [25], synchronized CL2006 or AGD1246 worms were grown on NG plates containing 100μg/ml ampicillin, spotted with E. coli cultures that express dsRNA as indicated. On day one of adulthood, 120 worms were transferred onto 10 5mm NG-ampicillin plates (12 animals per plate). These 10 plates were randomly divided into 5 sets (2 plates, 24 worms per set). Paralysis of these worms was scored daily by gently tapping their noses with a platinum wire or placing a hot pick proximally to their bodies. Worms that were capable of moving their noses but unable to move the trunk of their bodies were scored as “paralyzed” and removed from the plates. The assay was terminated at day 12 or 13 of adulthood in order to avoid scoring old animals as paralyzed. As a control, this assay was also performed using wild type N_2_ worms.

To follow the toxicity of polyQ35-YFP stretches by the “thrashing assay” [37], synchronized eggs of AM140 or AM1126 worms were placed on plates which were seeded with control bacteria (EV) or bacteria that express RNAi towards *skn-1*. At the indicated ages, one worm was placed in a drop of M9 buffer and the number of body bends per 30 seconds was scored. At each time point at least 20 animals were used. As a control, this assay was also performed using wild type N_2_ worms.

### Statistical analyses

To achieve statistical significance the Student T-test using two-tailed distribution and two-sample equal variance was used. The analyses of the experiments were conducted using at minimum of three independent biological repeats of each experiment as indicated. Statistical information of lifespan experiments is presented in the supplemental tables.

## Results

To examine when SKN-1 regulates lifespan in wild type worms (strain N_2_), we knocked down the expression of *skn-1* by RNAi at different stages during the worm’s lifecycle (Fig. S1A shows the high efficiency of the RNAi as tested using feminized CF512 worms, a strain whose lifespan is similar to that of wild type animals. We used these worms to avoid the possible effects of developing embryos on gene expression). Synchronized eggs of N_2_ animals were placed on plates seeded with control bacteria harboring the empty vector RNAi plasmid (EV), or with *skn-1* RNAi expressing bacteria. At larval stages L2, L4 or at day 1, 5 or 9 of adulthood, groups of 120 worms were picked from EV plates and transferred onto *skn-1* RNAi bacteria. Lifespans were followed by daily scoring of dead animals. While worms that were grown throughout life on EV bacteria had a mean lifespan of 18.12±0.51 days (±SEM), animals that were treated from hatching with *skn-1* RNAi exhibited a significantly (p<0.001) shorter mean lifespan of 16.17±0.32 days (Fig. 1A and B and Supplemental Table 1). This result is consistent with the previously reported shortening effect of *skn-1* mutation on the lifespan of these worms [12]. Interestingly, the knockdown of *skn-1* from the L2 or L4 larval stages resulted in similar lifespan shortening effects of 11.8% (mean lifespan 15.98±0.24 days, p<0.001) and 14.23% (mean lifespan 15.54±0.35 days, p<0.001), respectively (Fig. 1A and Supplemental Table 1). These lifespans were very similar to those of animals that were grown on *skn-1* RNAi throughout life (mean lifespan of 16.17±0.32 days, p<0.001). In contrast, the knockdown of *skn-1* from day 1 and 5 of adulthood resulted in a trend of lifespan shortening, however, this effect was not significant (Fig. 1B and Supplemental Table 1, mean lifespan 17.83±0.31 days, p=0.31 and mean lifespan 17.51±0.42 days, p=0.18, respectively). These results show that *skn-*1 is primarily needed later than the L4 larval developmental stage through early adulthood to regulate lifespan.

**Figure 1:**
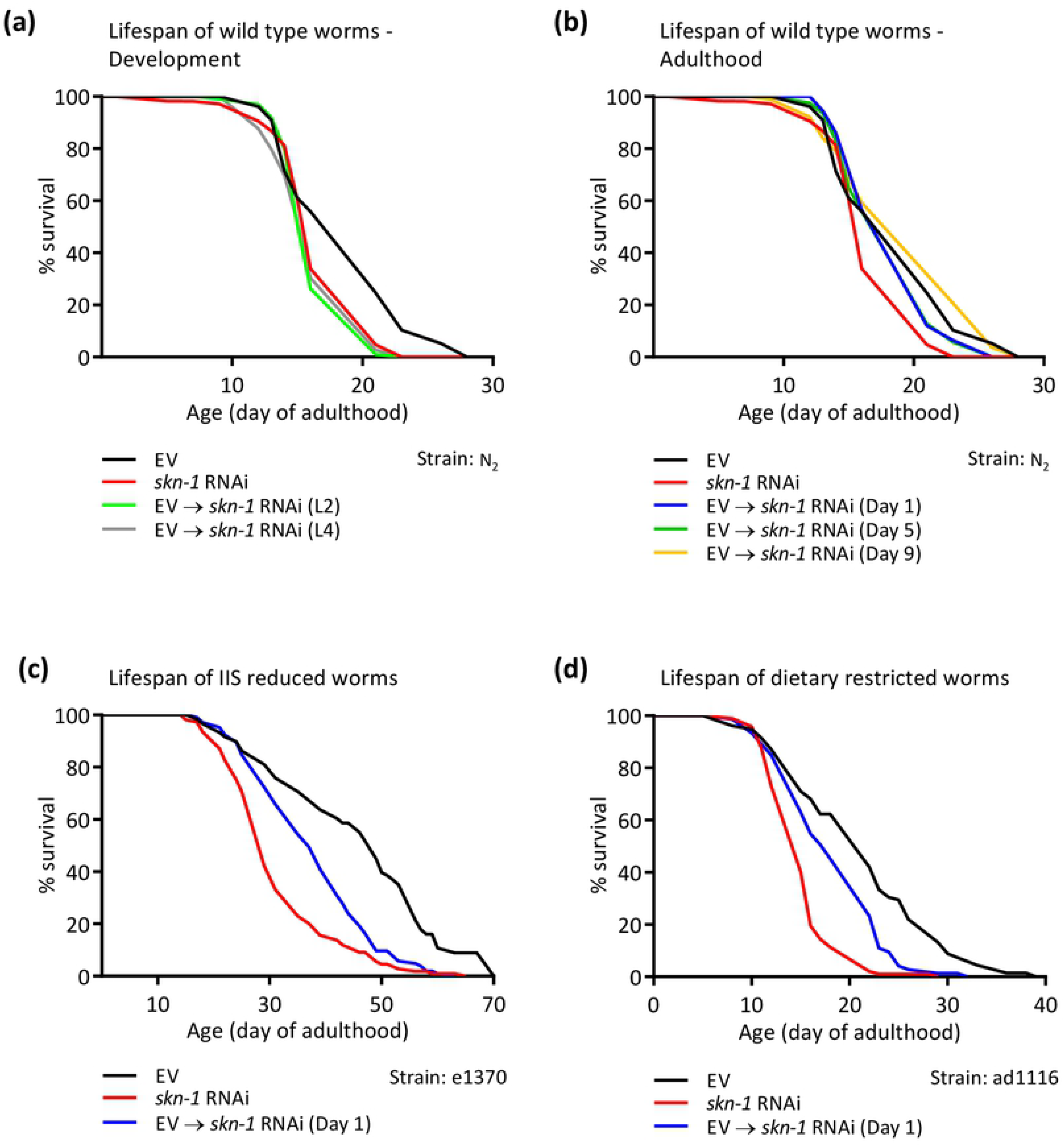
*skn-1* regulates lifespan primarily from late larval development through early adulthood. (**A-B**) The lifespan of wild type animals (WT, strain N_2_) treated from hatching with empty vector bacteria (EV, control), *skn-1* RNAi, or transferred from EV bacteria onto *skn-1* RNAi either during developmental stages L2 or L4 (A), or at day 1, 5, or 9 of adulthood (B) was measured. Worms treated with *skn-1* RNAi throughout life, or during developmental stages L2 or L4 showed significant reductions in lifespan (10.75%, 11.80%, and 14.23%, respectively, Supplemental Table 1). Worms treated with *skn-1* RNAi from day 1 or 5 of adulthood showed a trend of reduction in lifespan, though the observed lifespan shortening was not significant (Supplemental Table 1). Treating worms with *skn-1* RNAi from day 9 of adulthood did not affect lifespan (Supplemental Table 1). (**C-D**) The knockdown of *skn-1* throughout life or from day 1 of adulthood, resulted in lifespan shortening of *daf-2* (*e1370*) mutant worms (C and Supplemental Table 2A) and of *eat-2* (ad1116) mutant animals (D and Supplemental Table 3A).

To further examine the timing requirements of SKN-1 for longevity assurance, we employed long-lived mutant worm strains. *daf-2* (*e1370*) mutant worms carry a weak *daf-2* allele and thus, exhibit exceptional longevity [4]. The animals were grown throughout life either on EV or on *skn-1* RNAi bacteria. An identical group of worms was hatched on EV bacteria and transferred onto *skn-1* RNAi bacteria at day 1 of adulthood. While the knockdown of *skn-1* from hatching resulted in lifespan shortening of 31.6% compared to the lifespan of untreated worms (mean lifespan of 30.63±0.94 and 44.79±1.97 days, respectively), knocking down the expression of this gene exclusively during adulthood shortened lifespan by only 17.34% (Fig. 1C and Supplemental Table 2A, mean lifespan of 37.02±1.02). These results, which were verified with an additional biological experimental repeat (Supplemental Table 2B), indicate that *skn-1* is needed from day 1 of adulthood as a modulator of lifespan. Nevertheless, the observation that knocking down *skn-1* from day 1 of adulthood did not shorten lifespan as efficiently as *skn-1* RNAi treatment throughout life, suggests that this transcription factor is also needed during larval development to allow *daf-2* mutant worms to exhibit their full longevity potential. We repeated this experiment using ad1116 worms which carry a mutation in the *eat-2* gene, resulting in a pharyngeal defect that leads to constitutive dietary restriction, and thus, are long-lived [38]. We found that similarly to *skn-1* RNAi-treated *daf-2* (*e1370*) mutant worms, the knockdown of *skn-1* throughout life shortens the lifespan of ad1116 animals by 27.89% (mean lifespan of 14.93±0.35 days). The lifespan of their counterparts who were treated with *skn-1* RNAi from day 1 of adulthood was shortened by 13.58% (mean lifespan of 17.89±0.60 days), relative to the control worms (Fig. 1D and Supplemental Table 3A, mean lifespan 20.70±0.90 days). These results were confirmed with an additional biological experimental repeat (Supplemental Table 3B). Together, these results indicate that SKN-1 is at least partially required during development as a regulator of lifespan. However, SKN-1 is also needed during adulthood to promote the natural lifespan of wild type animals and confer the full longevity of long-lived mutant worms.

We next investigated when during the worm’s lifecycle SKN-1 regulates proteostasis. To determine this, we utilized CL2006 worms which express the AD-causing amyloid beta (Aβ) peptide in their body wall muscles [39]. This expression results in a progressive paralysis within the worm population; a phenotype that can be tracked by the “paralysis assay”, a daily scoring of paralyzed animals [25]. First, we tested whether the knockdown of *skn-1* throughout life enhances the paralysis phenotype of these animals and found that it does (Fig. 2A). The significance of this phenotype was established by three independent repeats of the paralysis assay (Fig. 2B and S1B). Importantly, the knockdown of *skn-1* in wild type worms did not enhance the rate of aging-associated paralysis up until day 12 of adulthood (Fig. S2A). To test whether the paralysis phenotype is tissue specific, we performed an identical experiment using AGD1246 worms which express the Aβ peptide under the regulation of the *rgef-1* pan-neuronal promoter [28]. We found, similarly to the observed phenotype in muscles, that the knockdown of *skn-1* by RNAi results in an increased rate of paralysis of these worms (Fig. S2B and C).

**Figure 2:**
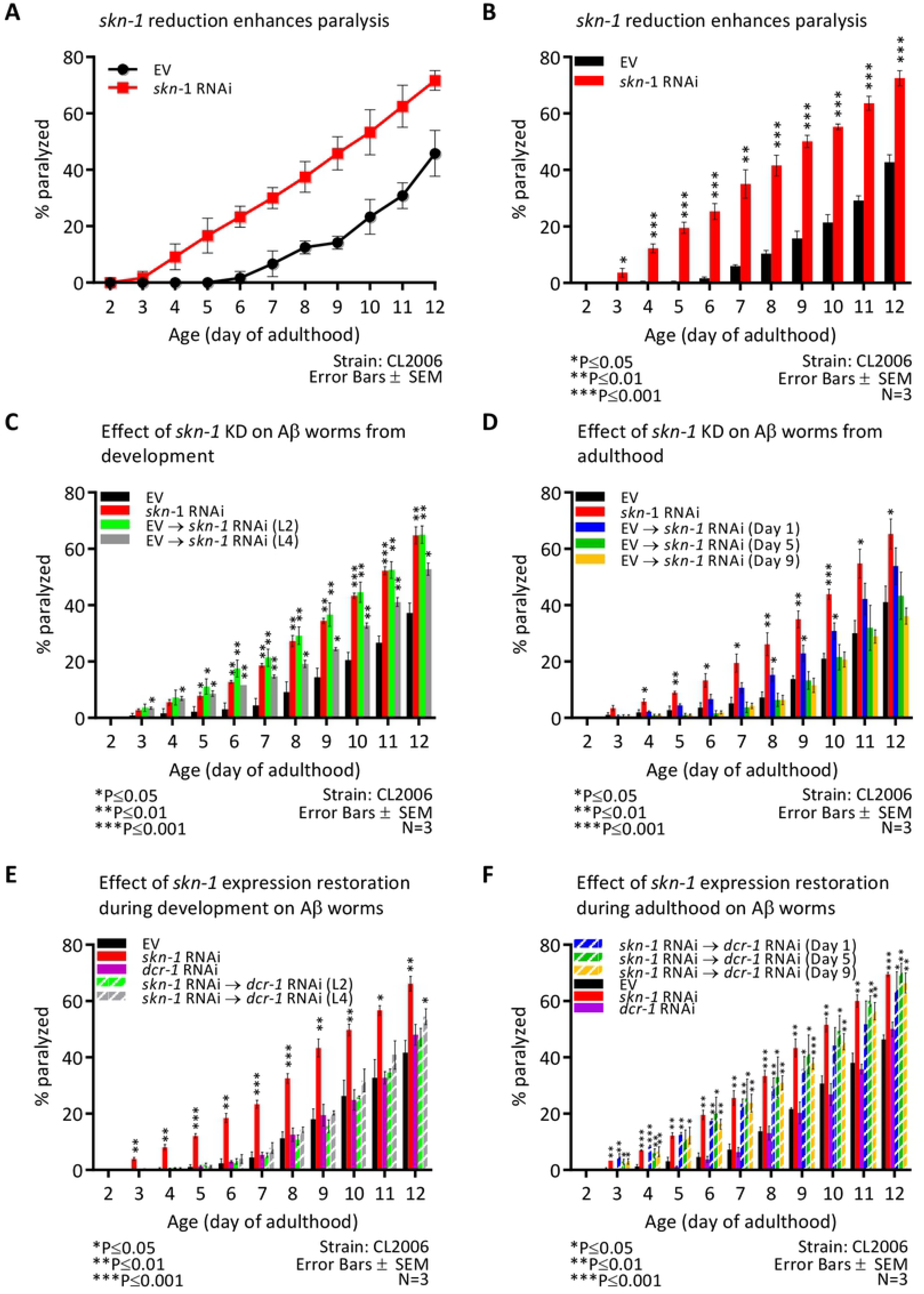
*skn-1* is foremost important from the L4 stage of larval development through day 1 of adulthood to protect against Aβ-induced proteotoxicity. (**A-B**) The knockdown of *skn-1* enhances the rate of paralysis of Aβ worms (A). Three independent repeats indicate that this effect is significant (B, p<0.001 see also Fig. S1B). (**C**) Treating Aβ worms with *skn-1* RNAi from the L2 larval stage enhances the rate of paralysis as efficiently as lifelong treatment with *skn-1* RNAi. In contrast, knocking down *skn-1* RNAi from the L4 larval stage less pronouncedly, but significantly enhances the rate of paralysis (see also Fig. S3A). (**D**) Worms grown from hatching on EV and transferred at day 1 of adulthood onto *skn-1* RNAi show a marginal increase in paralysis compared to worms grown for life on EV. In contrast, worms transferred onto *skn-1* RNAi at day 5 or 9 of adulthood show no increased rate of paralysis compared to their counterparts grown on EV throughout life (see also Fig. S3B). (**E**) Treating worms with *dcr-1* RNAi throughout life, or growing worms on *skn-1* RNAi from hatching and transferring them to *dcr-1* RNAi during the L2 or L4 stages of development did not enhance the rate of paralysis, apart from a very slight increase at day 12 of adulthood in the population transferred at the L4 stage (see also Fig. S3C). (**F**) Aβ worms that were grown on *skn-1* RNAi from hatching then transferred onto *dcr-1* RNAi at either day 1, 5 or 9 of adulthood and animals that were treated with *skn-1* RNAi throughout life, exhibited similarly enhanced rates of paralysis compared to the EV treated population (see also Fig. S3D).

To establish the temporal requirement of *skn-1* as a modulator of proteostasis we treated CL2006 worms with *skn-1* RNAi throughout the experiment or from the L2 or L4 larval stages. An identical group of CL2006 worms was grown throughout the experiment on EV bacteria. Three independent experiments indicated that worms which were treated with *skn-1* RNAi throughout the experiment and their counterparts that were transferred onto *skn-1* RNAi bacteria at the L2 developmental stage, were paralyzed at similar rates significantly higher than that of untreated animals (EV). These results indicate that *skn-1* plays no roles during early development (L2 stage and earlier) as a modulator of Aβ-mediated proteotoxicity. Worms that were transferred onto *skn-1* RNAi from the L4 larval stage exhibited a higher rate of paralysis than the control group (EV). However, the rate of paralysis of these worms was lower than that of nematodes that were treated from the L2 stage (Fig. 2C and S3A). This shows that SKN-1 activity in late developmental stages is needed for partial protection from Aβ.

To test whether *skn-1* is required during adulthood as a modulator of proteostasis, we conducted a similar experiment in which CL2006 worms were grown on EV bacteria and then transferred onto *skn-1* RNAi at either day 1, 5 or 9 of adulthood. Rates of paralysis were scored daily. Three independent experiments showed that the knockdown of *skn-1* at day 1 of adulthood enhances the rate of paralysis (Fig. 2D and S3B). This effect, however, was less prominent than that of knocking down *skn-1* during development (Fig. 2C and S3A). No significant enhancement in the paralysis phenotype, compared to untreated worms, was observed when the worms were treated with *skn-1* RNAi from day 5 or 9 of adulthood (Fig. 2D and S3B).

These results propose that SKN-1 is foremost required as a proteostasis regulator during late larval development through early reproductive adulthood. To further test this conclusion, we conducted a reciprocal set of experiments using *dcr-1* RNAi. DICER, encoded by *dcr-1*, is a nuclease that cleaves double stranded RNA to create small interfering RNA (siRNA) and thus, is crucial for the functionality of the RNAi machinery [40]. Accordingly, the knockdown of *dcr-1* by RNAi inactivates the RNAi machinery and restores the expression of the knocked down gene to near natural levels [33]. We utilized this technique to conditionally knockdown *skn-1* and followed the rates of paralysis of CL2006 worm populations that hatched on *skn-1* RNAi bacteria, and were then transferred onto plates seeded with *dcr-1* RNAi at the L2 or L4 larval stages. Three experimental repeats indicated that the knockdown of *dcr-1* had no effect on the rate of paralysis, as animals that were grown on control bacteria (EV) and their counterparts that were treated with *dcr-1* RNAi throughout the experiment, had similar rates of paralysis (Fig. 2E and S3C). As expected, the knockdown of *skn-1* throughout the assay increased paralysis. However, knocking down *skn-1* solely during early development, from hatching up until the L2 larval stage, did not increase the rate of paralysis. The knockdown of *skn-1* from hatching up until the L4 larval stage had a small deleterious effect, as the rate of paralysis was significantly higher than that of untreated animals solely at day 12 of adulthood. These results suggest that SKN-1 is needed as a regulator of proteostasis from the L4 stage of larval development.

To further scrutinize the temporal requirements of *skn-1* as a regulator of proteostasis, we tested how the knockdown of *skn-1* affects the paralysis of Aβ worms during adulthood. Synchronized eggs were placed on plates that were seeded with *skn-1* RNAi bacteria and transferred onto *dcr-1* RNAi plates on either day 1, 5 or 9 of adulthood. Our results (Fig. 2F and S3D) indicate that worms treated with *skn-1* RNAi throughout development and transferred onto *dcr-*1 RNAi at either day 1, 5 or 9 of adulthood, exhibited similar rates of paralysis to animals fed with *skn-1* RNAi bacteria throughout life.

Together our results demonstrate that SKN-1 is required for protection against Aβ induced proteotoxicity from the L4 stage of larval development through the first day of adulthood. However, the restoration of *skn-1* expression at day 5 or 9 of adulthood did not rescue the enhanced paralysis phenotype, indicating that this transcription factor is dispensable as a proteostasis regulator in late stages of adulthood. These temporal requirements show that SKN-1 is required subsequently to HSF-1, which is primarily required during the L2 larval developmental stage [34], for proteostasis maintenance..

We next sought to test whether this temporal pattern of SKN-1 as a proteostasis modulator is also true for worms that are challenged by the aggregation of a proteotoxic protein other than Aβ. To address this, we utilized worms that express a chimeric, fluorescently-tagged polyglutamine protein of 35 repeats (polyQ35-YFP) in their body wall muscles (strain AM140). Abnormally long polyglutamine stretches in different proteins underlie the development of several human neurodegenerative maladies, including HD [22] and Machado Joseph Disease (MJD) [41]. AM140 animals accumulate aggregates and exhibit progressive motility impairment, [26], a phenotype that can be followed by the “thrashing assay” [37]. First, we tested whether the knockdown of *skn-1* affects polyQ35-YFP toxicity by comparing the thrashing rates of AM140 worms that were treated from hatching with *skn-*1 RNAi to those of untreated animals (EV). We found that the knockdown of *skn-1* results in a significantly reduced rate of motility on both day 2 and 6 of adulthood (Fig. 3A and B), while knocking down *skn-1* in wild type worms results in only a slight reduction in motility (Fig. S4A). Similar results to those observed in the AM140 worms were obtained when thrashing experiments were conducted using worms that express polyQ35-YFP under the *rgef-1* pan-neuronal promoter (strain AM1126, Fig. S4B and C), indicating that this phenotype is not tissue specific. We next tested when SKN-1 protects worms from polyQ35-YFP by growing AM140 animals on EV bacteria and transferring them onto plates that were seeded with *skn-1* RNAi bacteria at the L2 or L4 larval stages or at day 1 or 3 of adulthood. Thrashing rates were scored at days 3 and 6 of adulthood. Our results indicate that analogously to its roles in the mitigation of Aβ proteotoxicity, SKN-1 is foremost important as a regulator of proteostasis during late larval development through early stages of adulthood (Fig. 3C and D).

**Figure 3:**
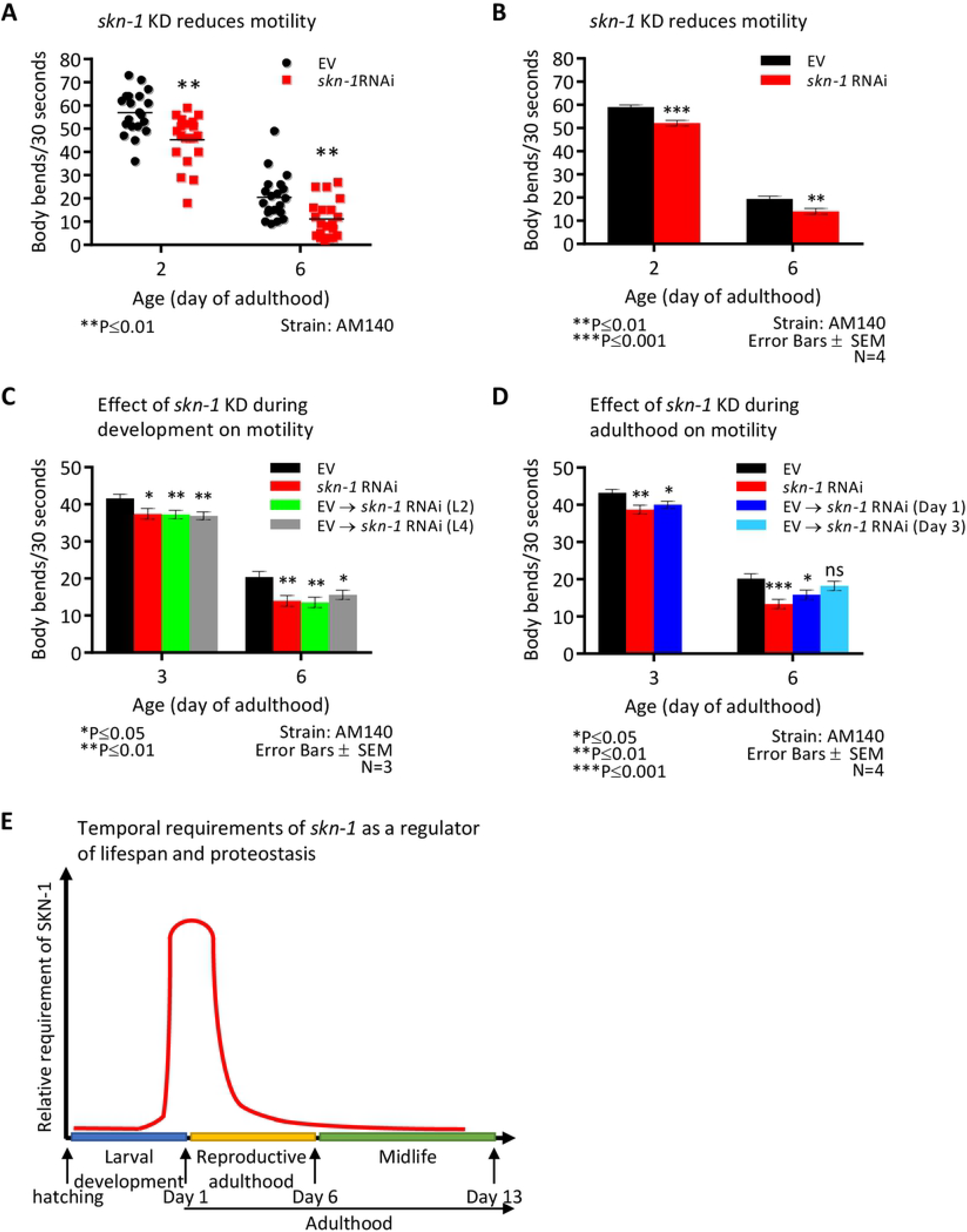
*skn-1* expression is most critical from the L4 larval stage through day 1 of adulthood to counter polyglutamine (polyQ) induced proteotoxicity. (**A**) The knockdown of *skn-1* by RNAi significantly decreased the thrashing rates of worms that express polyQ35-YFP in their muscles (strain AM140) at day 2 and 6 of adulthood. (**B**) Four independent repeats show that this effect is significant (reduction of 11.65% and 27.64%, respectively). (**C**) Growing AM140 worms on control bacteria from hatching followed by transferring them onto *skn-1* RNAi at the L2 or L4 developmental stage resulted in enhanced proteotoxicity as measured by the thrashing assay (Reduction of 10.45% and 11.29% at day 3, respectively, and a reduction of 33.63% and 23.59% at day 6, respectively). (**D**) Similarly, the knockdown of *skn-1* from day 1 (reduction of 10.38% and 7.35% when measured at day 3 and day 6, respectively), but not from day 3 of adulthood, resulted in a significant reduction in motility. (**E**) A schematic illustration of the temporal requirements for SKN-1 as a regulator of lifespan and proteostasis.

## Discussion

Our temporal analysis indicates that *skn-1* is predominantly required for lifespan determination and for protection from proteotoxicity, from the L4 stage of larval development through early adulthood (Fig. 3E). Because the IIS regulates lifespan [33] and proteostasis [35] during adulthood, it is likely that during the late stages of larval development through the early stages of adulthood, SKN-1 regulates the expression of gene networks that enable IIS reduction to promote these functions later in life. SKN-1 is required as a lifespan and proteostasis regulator in a time window which is subsequent to that of HSF-1, and partially overlapping and preceding the time window in which DAF-16 executes these functions [33–35]. Interestingly, SKN-1 is also at least partially needed during development for DR-promoted longevity (Fig. 1D). In contrast, the transcription factor PHA-4, which is also crucial for DR-mediated longevity, is solely needed during adulthood to enable this phenotype [42].

These observations substantiate that different transcription factors are needed in a sequential manner during the nematode’s lifecycle and raise the question of how SKN-1 acts, and what genes it regulates during late stages of development through early adulthood to enable the promotion of longevity and proteostasis in later stages of life. One possibility suggests that the regulation of stress responses, such as oxidative stress [43], by SKN-1, reduces damage during early stages of life. Accordingly, the knockdown of *skn-1* during these early stages, in which the organism may be more vulnerable to metabolic insults, results in a less efficient activation of stress response mechanisms, higher rates of damage accumulation, and accelerates the process of aging.

An alternative model suggests that during late larval development through early adulthood, SKN-1 regulates the expression of genes whose products are needed for IIS reduction and DR to promote lifespan and proteostasis in adulthood. This theme may be supported by the observation that *skn-1* is highly expressed during the L2 larval stage, a stage preceding the time window in which SKN-1 is most critical for lifespan and proteostasis regulation, compared to the observed expression during adulthood of CF512 worms (Fig. S5A). It is important to note that CF512 worms were shown to exhibit the same temporal requirements of *skn-1* expression (Fig. S5B and Supplemental Tables 4A and 4B) as seen in the lifespan experiments of wild type as well as long-lived worms (Fig. 1A-D). It would be interesting to compare the gene networks that are differentially regulated by SKN-1 in late larval development through early adulthood compared to other stages of life. Such target genes might encode constitutive heat shock proteins and inducible protective proteins. It would also be interesting to test whether SKN-1 and HSF-1 co-regulate target genes during development and whether the products of these genes are needed for the IIS to promote longevity and/or proteostasis during adulthood.

Another key question is where SKN-1 executes its longevity and proteostasis-promoting functions. Together, the known roles of neurons in the regulation of proteostasis [44, 45], the prominent expression of *skn-1* in ASI neurons [36] and the differential regulation of DAF-16 and HSF-1 by a neuronal gene [46], suggest that the developmental functions of SKN-1 may be regulated at the organismal level by neurons. It would also be interesting to test whether the activity of signaling complexes that reside on caveolae, a membrane structures that we previously found to regulate Aβ-mediated proteotoxicity [47], is affected by the knockdown of *skn-1.* Further research is needed to test these possibilities.

An additional important aspect of the temporal analyses of IIS regulated transcription factors is the tight correlations between longevity and proteostasis. While DAF-16 regulates both lifespan and proteostasis during adulthood [33, 35], and HSF-1 primarily during the L2 stage of larval development [34, 35], SKN-1 govern these functions primarily from the L4 stage of larval development through early adulthood. This correlation supports the notion that the formation of an efficient proteostasis assurance mechanism is needed for IIS reduction, and perhaps also for DR, to slow the progression of aging and promote longevity [48].

The requirement of *skn-1* during early adulthood as a regulator of lifespan and proteostasis overlaps with the reproductive adulthood time window in which *daf-16* is needed to enable longevity of *daf-2* mutant worms [33]. It is tempting to speculate that DAF-16 and SKN-1 may co-regulate the expression of certain genes during reproductive adulthood. Indeed, both SKN-1 and DAF-16 were shown to regulate the mitophagy mediator *dct-1* to regulate mitochondrial health [49]. This theme is also supported by the observation that the IIS and proteostasis-maintaining signaling that originate from the reproductive system are linked at the post-translational level [50] and by the recently reported integration of signals that originate from the reproductive system and from the IIS [51].

## Acknowledgments

This study was generously supported by the Israel Science Foundation (ISF) (EC#981/16), the Israeli Ministry of Science and Technology (MOST#80884) and the Henri J. and Erna D. Leir Chair for Research in Neurodegenerative Diseases.

## Author contributions

EC and DG initiated this project. DG performed most of the experimental work. AL performed lifespan assays and HB performed lifespan and proteotoxicity assays. EC wrote the manuscript.

## Competing interests

The authors declare no competing interests.

